# Effects of Soluble Electron Shuttles on Microbial Iron Reduction and Methanogenesis

**DOI:** 10.1101/2024.11.07.622564

**Authors:** Bhim Sen Thapa, Theodore M. Flynn, Zena D. Jensvold, Kenneth M. Kemner, Margaret F. Sladek, Edward J. O’Loughlin, Christopher W. Marshall

## Abstract

In many aquatic and terrestrial ecosystems, iron (Fe) reduction by microorganisms is a key part of biogeochemical cycling and energy flux. The presence of redox active electron shuttles in the environment potentially enables a phylogenetically diverse group of microbes to use insoluble iron as a terminal electron acceptor. We investigated the impact that different electron shuttles had on respiration, microbial physiology, and microbial ecology. We tested eight different electron shuttles, seven quinones and riboflavin, with redox potentials between 0.217 V and -0.340 V. Fe(III) reduction coupled to acetate oxidation was observed with all shuttles. Once Fe(III) reduction began to plateau, a rapid increase in acetate consumption was observed and coincided with the onset of methane production, except in the incubations with the shuttle 9,10-anthraquinone-2-carboxylic acid (AQC). The rates of iron reduction, acetate consumption, methanogenesis, and the microbial communities varied significantly across the different shuttles independent of redox potential. In general, shuttles appeared to reduce the overall diversity of the community compared to no shuttle controls, but certain shuttles were exceptions to this trend. Geobacteraceae were the predominant taxonomic family in all enrichments except in the presence of AQC or 1,2,-dihydroxyanthraquinone (AQZ), but each shuttle enriched a unique community significantly different from the no shuttle control conditions. This suggests that the presence of different redox-active electron shuttles can have a large influence on the microbial ecology and total carbon flux in the environment.

**Importance:** Iron is the fourth most abundant element in the Earth’s crust and the reduction of iron by microbes is an important component of global biogeochemical cycles. A phylogenetically diverse group of microbes are capable of conserving energy with oxidized iron as a terminal electron acceptor, but the environmental conditions favoring certain taxonomic clades in iron-reducing environments is unclear. One complicating factor often overlooked in small-scale enrichments is the influence of soluble, redox-active electron shuttles on the rate and microbial ecology of iron reduction. We tested the effects of eight different electron shuttles on the microbial physiology and ecology in iron-reducing enrichments derived from a local wetland. Each electron shuttle varied the microbial activity and enriched for a microbial community distinct from the no shuttle control condition. Therefore, in complex subsurface environments with many redox-active compounds present, we propose electron shuttles as a reason for the coexistence of multiple clades of iron-reducing bacteria.

## INTRODUCTION

Iron reduction by microorganisms is a significant component of biogeochemical cycling and energy flux in many aquatic and terrestrial environments. Dissimilatory metal-reducing bacteria (DMRB) are phylogenetically diverse microorganisms that obtain energy by coupling the oxidation of organic compounds or hydrogen to the reduction of iron and other metal oxides (1). As a group, DMRB can use a wide range of Fe(III) forms as terminal electron acceptors for anaerobic respiration including soluble Fe(III) complexes, Fe(III) oxides, and clay minerals containing varying amounts of structural Fe(III) (2–4). Because of the relative insolubility of most Fe-bearing minerals, their use by DMRB as terminal electron acceptors for respiration requires different mechanisms for electron transfer relative to soluble terminal electron acceptors that are easily transported into the cell (e.g., O_2_, NO_x_ ^-^, and SO_4_ ^2-^) (5). One approach involves the transfer of electrons from the cell to external electron acceptors by soluble electron shuttles (e.g., quinones, flavins, phenazines, and reduced sulfur species) (6–8).

Electron shuttles (or shuttles) are molecules that can reversibly donate or accept electrons, thereby acting as oxidants or reductants in redox reactions. Shuttles containing quinone groups, due to electron delocalization among conjugate bonds, act as excellent electron mediators in biological energy metabolism (9), and are key players in redox processes in aquatic and terrestrial environments (10). Indeed, the use of quinone groups within humic substances (a major component of natural organic matter (NOM) consisting of heterogeneous mixtures of polydispersed materials resulting from the decay and transformation of plant and microbial remains) as electron shuttles is an important pathway for microbial reduction of Fe(III) oxides in anoxic water/soils/sediments (11–13). In addition to utilizing exogenous shuttles, many microbes can synthesize electron shuttling compounds, including flavins, nicotinamide adenine dinucleotide (NAD), membrane bound quinones, cytochromes, and phenazines (8, 14–18).

Dissimilatory reduction of Fe(III)-bearing minerals in anoxic environments is commonly observed in association with *Geobacter* species (19), and a large body of literature exists describing the mechanisms of direct electron transfer (shuttle-less) to Fe(III) oxides by outer membrane cytochromes in *Geobacter* spp. (18, 20–22). *Shewanella* spp. are also well-studied metal-reducing bacteria found in aquatic and terrestrial environments that can directly (via cytochromes) or indirectly transfer electrons to extracellular electron acceptors, most notably through production of flavin shuttles (14, 15, 23–25). Moreover, some microbes can only access insoluble terminal electron acceptors using electron shuttles (26). Because of the varied iron-reducing capacity of different microbial taxa, we hypothesized electron shuttles would increase the diversity of microbial communities under Fe(III)-reducing conditions by allowing greater access to insoluble terminal electron acceptors as the presence of electron shuttles can allow for the reduction of Fe(III) oxides by organisms that are not generally recognized as Fe(III)-reducing microorganisms (e.g., methanogens, sulfate reducers, and fermenters) (8, 27–32). We also hypothesized shuttles would increase the rate of iron reduction in microcosm enrichments. To test how electron shuttles influence microbial ecology and metabolic activity, we established enrichments with different electron shuttles covering a range of redox potentials (Table 1) and measured acetate consumption, Fe(III) reduction, CO_2_ production, methanogenesis, and microbial community changes.

**Table 1.**
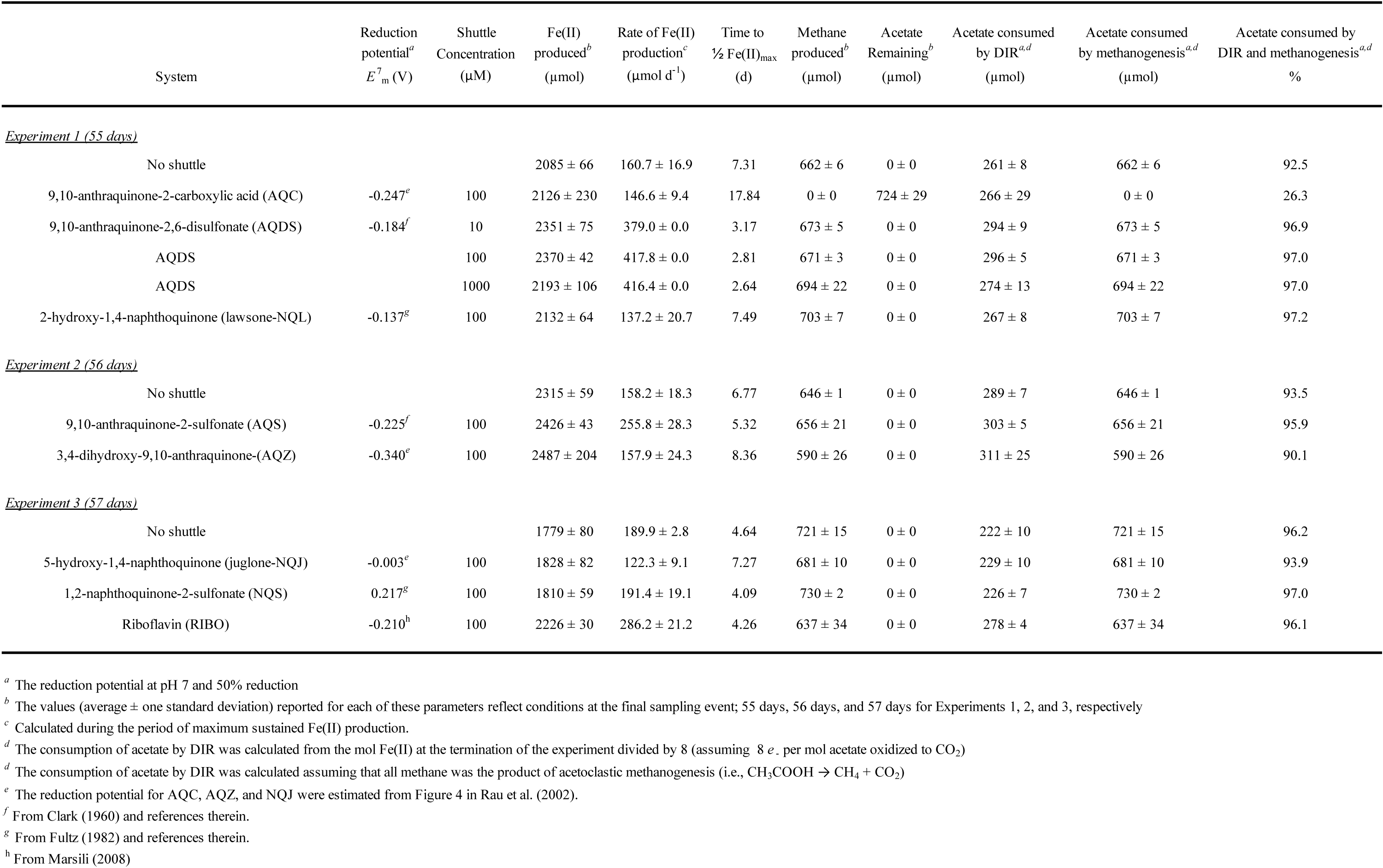
Electron shuttle concentration and reduction potential, total Fe(II) production, rate of Fe(II) production, time needed to reach one half maximum Fe(II) concentration, methane production, residual acetate, acetate consumed by DIR, acetate consumed by methanogenesis, percent of acetate consumed by both DIR and methanogenesis.

## RESULTS

### Effects of shuttles on carbon and electron flow

To understand the effects that environmentally relevant, soluble electron shuttles have on iron reduction and methanogenesis, we established replicate enrichments that were inoculated with a sediment slurry derived from a freshwater wetland (referred to as ’original inoculum’). We incubated the enrichments anaerobically with one of eight different shuttles or the no shuttle control. Three different sets of replicate no shuttle control enrichments were run with each batch of experiments (n=9) to ensure that we could make meaningful comparisons across our experiments. We observed only a small amount of variation in the measured metabolic parameters (acetate consumption, iron reduction, and methane generation) between replicates within and between experimental batches in the no shuttle controls (Figure 1, gray box and whisker plots), indicating that our results were reproducible and comparable between experimental batches.

**Figure 1.**
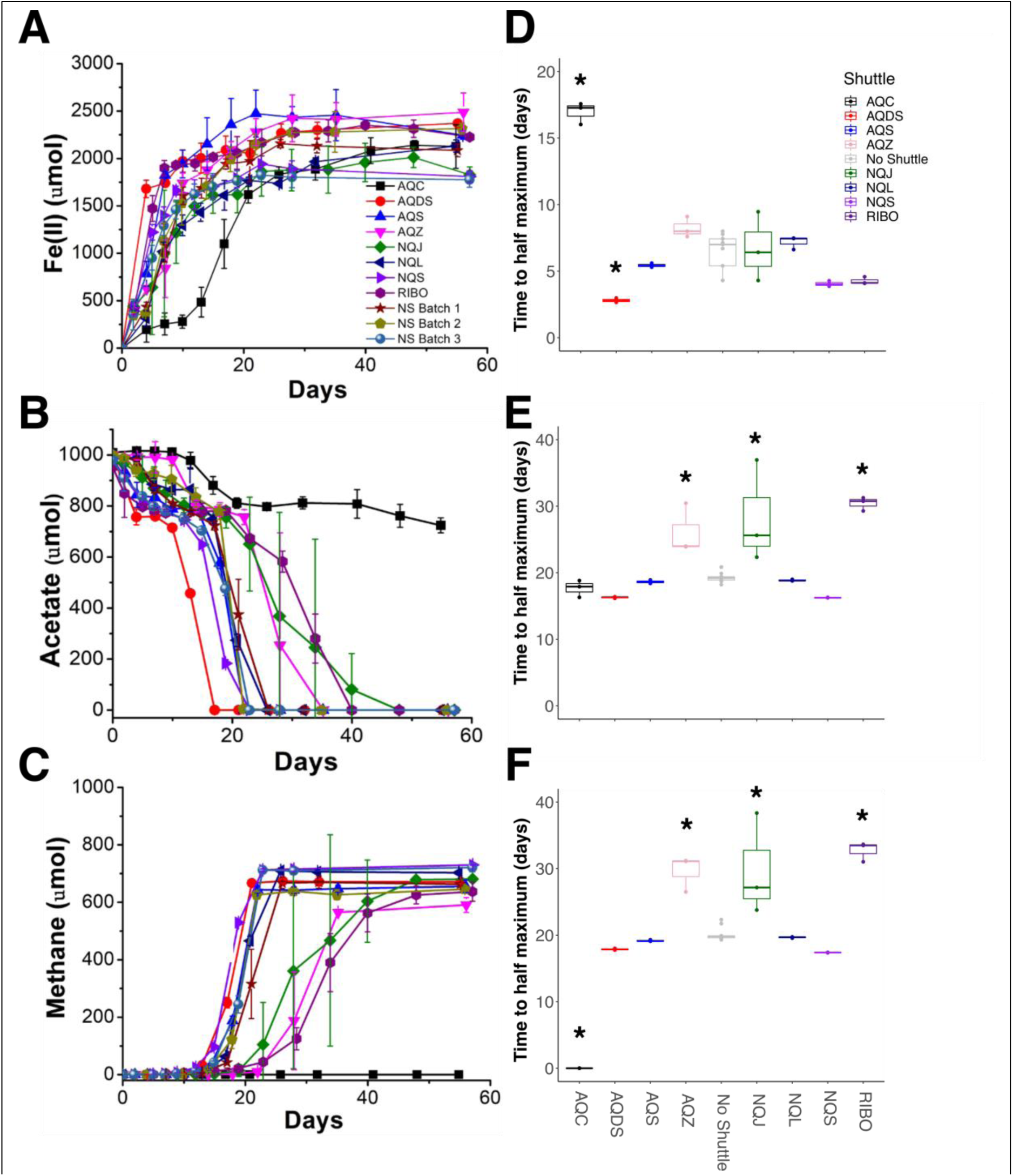
Iron, acetate, and methane reduction or oxidation rates for each shuttle over the course of the experiments. (A) Fe(II) production, (B) acetate consumption, (C) methane production for each shuttle over time. (D-F) Rates of substrate reduction or oxidation measured as time to half the maximum concentration. Shuttles AQC black squares, AQDS red circles, AQS blue up triangles, AQZ pink down triangle, No shuttle batch 1 maroon star, No shuttle batch 2 tan pentagon, No shuttle batch 3 grey-blue sphere, NQJ black left triangle, NQL brown right triangle, NQS dark blue octagon, and Riboflavin purple star. Asterisks over the shuttle rate values indicates significantly faster or slower rates compared to the no shuttle controls, p<0.05, ANOVA with Tukey’s multiple comparison correction. All experiments n=3, except in right panels where no shuttle batches were aggregated (gray), n=9.

Regardless of the particular shuttle present, Fe(III) reduction coupled to acetate consumption was observed in all of the enrichments (Figure 1). During the first two weeks, acetate consumption was similar across all shuttle treatments and corresponded to a reduction of ∼10% of the available Fe(III), with the exception of the AQC shuttle treatment (Supplemental Figure S1). Iron reduction rates were shuttle dependent, where the presence of AQDS significantly increased iron reduction compared to the no shuttle control and AQC significantly slowed iron reduction rates compared to the no shuttle control (Figure 1D). Following this initial phase where acetate and Fe(II) concentrations appeared to plateau, a rapid rise in acetate oxidation coupled to an increase in methane production was observed (Figures 1B-C). This second phase, where only minimal iron reduction was observed, was more variable by treatment and depended on which shuttle was present (Supplemental Figure S1). In particular, there was no significant difference in the rate of acetate consumption between AQS, AQDS, NQL, NQS, or no shuttle (NS), but the presence of shuttles AQC, AQZ, NQJ and riboflavin impaired the acetate consumption and methanogenesis rates (Figure 1E-F). The enrichments containing AQC behaved uniquely compared to the rest of the shuttles, with a significantly lower amount of acetate consumption, and a complete inhibition of methanogenesis (Figure 1). At the conclusion of the experiment, the total amount of acetate consumed and accounted for in dissimilatory iron reduction and methanogenesis, with the exception of AQC, was between 900 and 970 µM, corresponding to 90–97% of the entire amount of acetate that was available (Table 1). Therefore, the particular shuttle had a significant effect on the rate of substrate utilization, but the total amount of substrate consumed did not differ at the end of the experiment, except with AQC present. In addition, this reinforces the methodology of the current work, where the total concentration of acetate provided an upper limit for product formation and very little substrate was from the original inoculum.

### Microbial Community Dynamics

Given the variable rates of acetate consumption, Fe(II) production, and methanogenesis based on electron shuttle addition, we predicted that the shuttles also altered the microbial ecology of the system. The overall within-sample diversity of the enrichments, measured using the Shannon index, significantly varied by shuttle treatment (Figure 2). Unsurprisingly, the original inoculum (three different freshwater wetland sediment slurries prepared at the beginning of each batch experiment) had significantly more diverse microbial communities compared to the subsequent enrichments. Contrary to our initial hypothesis that the addition of shuttles in general would increase diversity under Fe(III)-reducing conditions, the no shuttle control enrichments had the highest diversity and was similar to AQDS, NQL, NQS, and riboflavin. The enrichments containing AQC, AQS, and AQZ had the lowes t divers ity.

**Figure 2.**
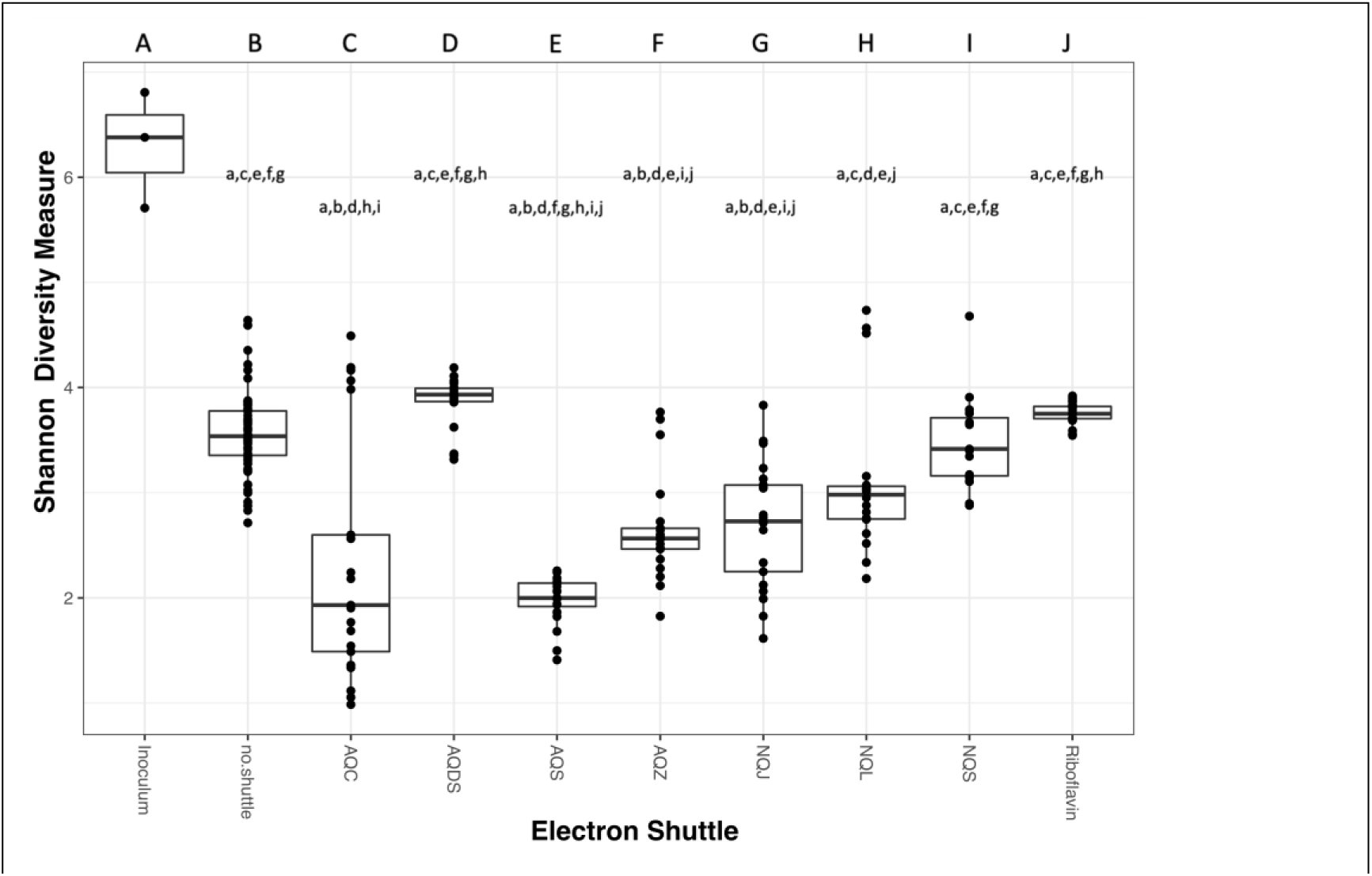
Alpha diversity across all shuttle treatments as measured by the Shannon diversity index. Each column is assigned an upper-case letter and if a treatment has that letter in lower-case that means they are significantly different, p<0.05, ANOVA with Tukey’s multiple comparison correction.

In general, taxa from the Geobacteraceae family were the predominant taxa across all treatments (Figure 3). However, based on the Bray-Curtis dissimilarity measure of between sample diversity, each electron shuttle treatment had a distinct microbial community from the no shuttle treatment (Figure 4) and the relative abundance of Geobacteraceae varied between shuttle treatments (Figure 5). The highest relative abundance of Geobacteraceae was observed at 70-80% of the community in AQS and NQL followed by 40-60% of the community in NQJ and NQS treatments. Of the genera within Geobacteraceae that could be assigned, *Citrifermentans*^1^ was the predominant genus across AQDS, AQS, NQL, and NQS whereas the *Geobacter* genus was predominant in riboflavin (Figure 5A).

**Figure 3.**
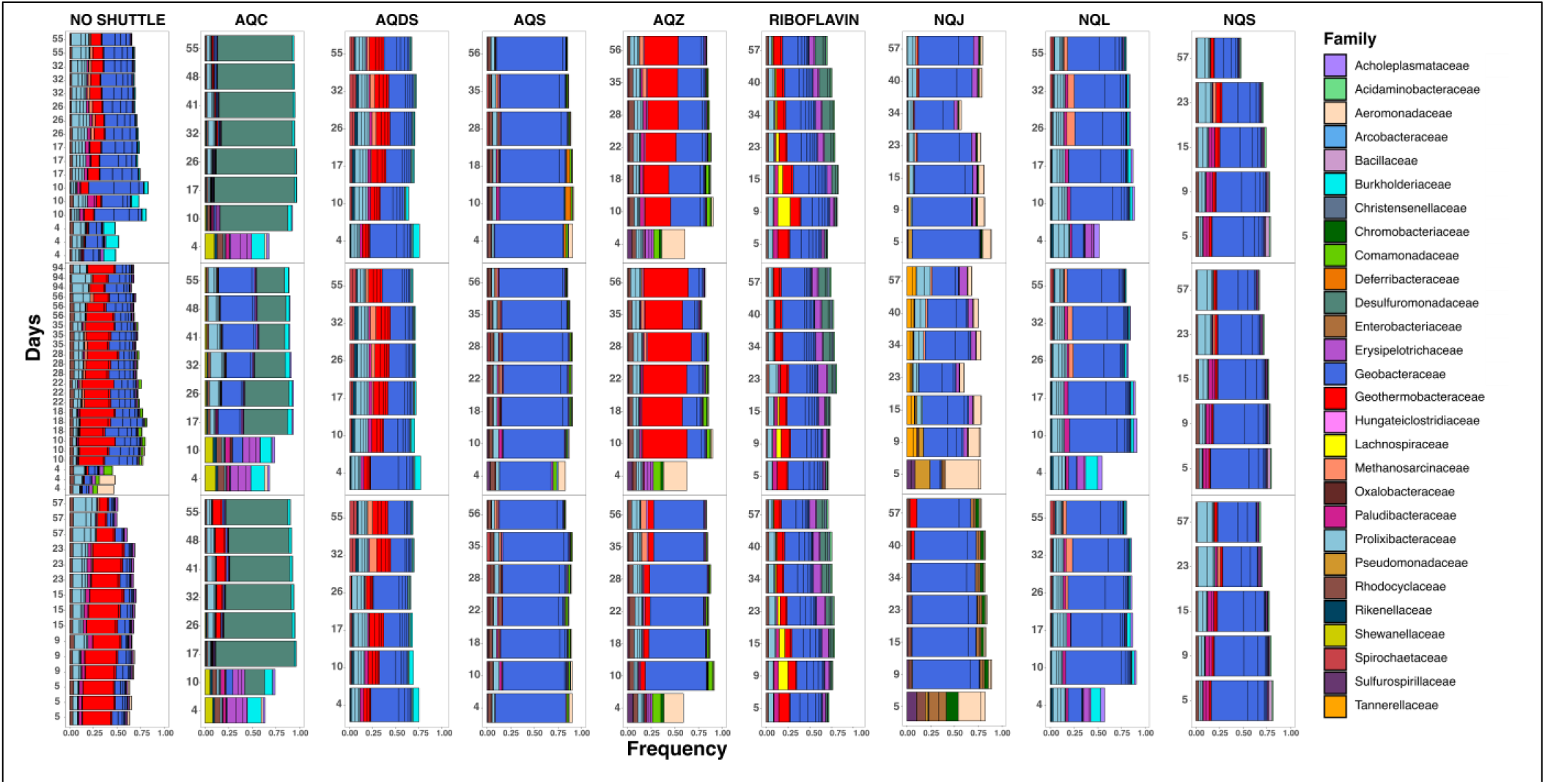
Stacked bar plots of the top 20 most abundant taxonomic families in the no shuttle treatment and in the shuttle treatments. The no shuttle treatment has plots for batch1, batch2, and batch3 from top to bottom and each plot contain three replicates each whereas the shuttle treatments represent the three replicates in the three different panels from top to bottom.

**Figure 4.**
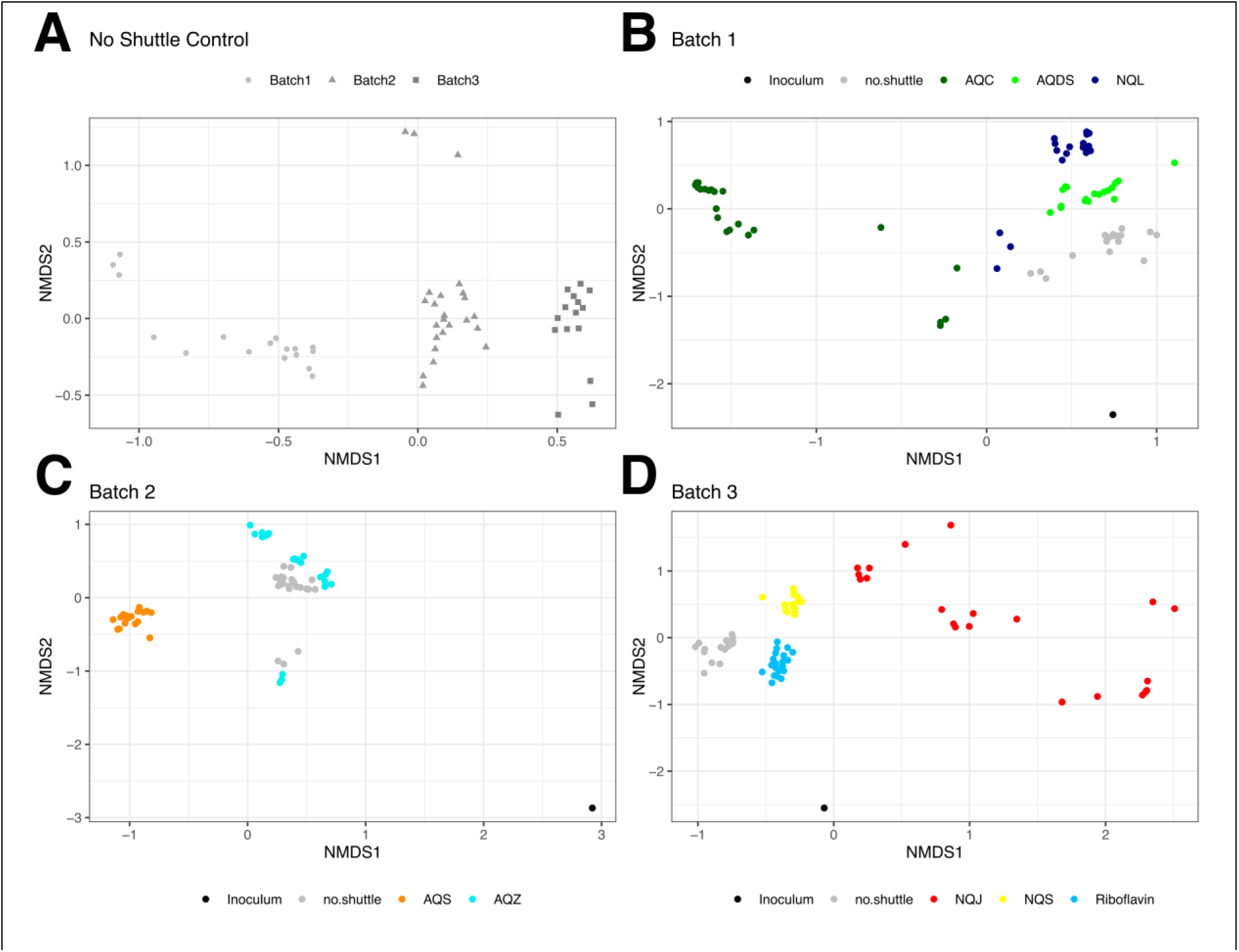
Non-metric multidimensional scaling (NMDS) ordination plots using the Bray-Curtis distance between communities. The no shuttle treatments are compared between experimental batches (A) and then each shuttle is compared to the no shuttle treatment associated with the appropriate experimental batch in (B-D).

**Figure 5.**
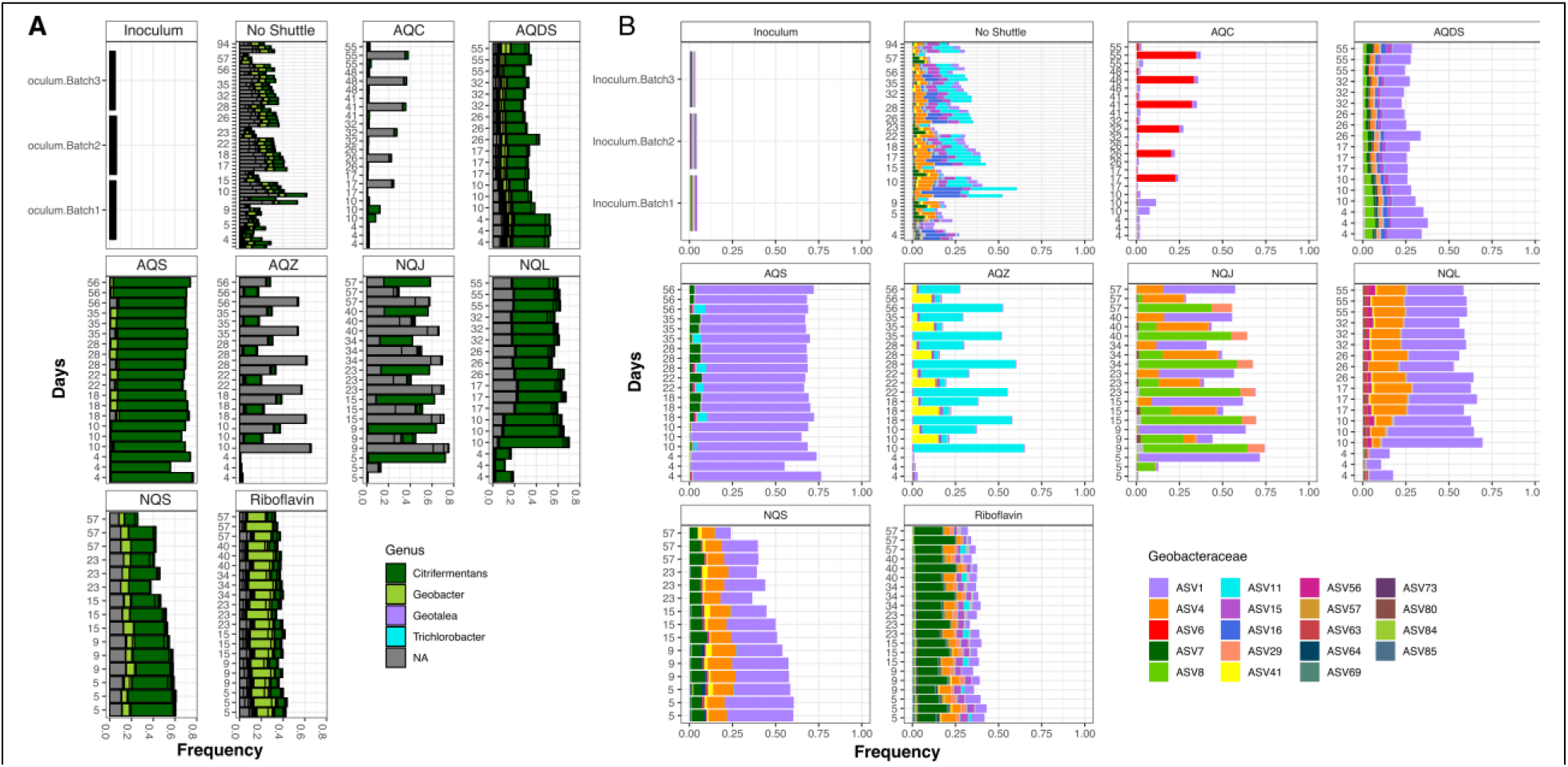
Stacked bar plots of the genera within the Geobacteraceae family for each shuttle treatment (A) and amplicon sequence variants (ASV) of *Geobacteraceae* family observed in different shuttles (B). N/A indicates not assigned at the genus level. The plot for each shuttle contains all three replicates per shuttle.

A small group of amplicon sequence variants (ASVs) belonging to the Geobacteraceae family accounted for over 50% of the microbial community in several treatments (Figure 5B). These ASVs provide insight into how each shuttle might select for specific taxa within the same family. AQS, NQS, NQL, and one of the replicates of NQJ had high relative abundance of ASV1, classified as *Citrifermentans*. Unique in the high relative abundance to NQJ was ASV8, classified as *Geobacter*, and in high relative abundance in AQZ was ASV11, classified as a different *Geobacter* variant (Figure 5A).

Beyond Geobacteraceae, other enriched taxa appeared to be shuttle treatment-specific (Figure 3). The Geothermobacteraceae family was one of the top taxonomic assignments in the no shuttle control, but only AQZ shuttle enriched for high abundance of that family. Taxa from the Desulfuromonadaceae family were by far the predominant taxa enriched in the AQC enrichment, likely reflecting the unique pattern of iron reduction and acetate consumption observed in this shuttle treatment.

Taxa assigned to the Halobacterota phylum, primarily *Methanosarcina*, reached 1% relative abundance in many of the treatments only after day 15, after which the abundance began to increase (Supplemental Figure S3), concurrent with the onset of methane production (Figure 1). Consistent with the lack of methanogenesis in the AQC shuttle treatment, Halobacterota were not observed in the presence of that shuttle.

As these experiments were conducted in three different batches with inoculum collected from the same location but at different times, we compared the no shuttle control condition across each batch. Non-metric multidimensional scaling (NMDS) was used to visualize the differences in the microbial communities (using the Bray-Curtis dissimilarity measure) between treatments (Figure 4, Supplemental Figure S2). Because the no shuttle controls were significantly different between batches (Figure 4A), we compared each shuttle community to the community of the no shuttle treatment run in the same batch of experiments. Each shuttle microbial community was significantly different from the associated no shuttle treatment (PERMANOVA with Benjamini and Hochberg adjustment, 10,000 permutations, p<0.05). This demonstrates that each electron shuttle exerts a significant and measurable influence on the overall microbial community.

## DISCUSSION

Our data show that the availability of different electron shuttles can significantly alter carbon metabolism, Fe(II) production rates, and microbial community composition in sediment-based enrichments. Since quinone-based shuttles can be used as electron acceptors by taxonomically diverse bacteria (36–38), we hypothesized that the presence of electron shuttles would increase the overall microbial community diversity under Fe(III)-reducing conditions by increasing terminal electron acceptor availability. Instead, our results indicate that diversity decreased in the presence of several of the shuttles as indicated by the lower alpha diversity in AQC, AQS, AQZ, and NQJ compared to the no shuttle treatment (Figure 2). However, while the presence of a single electron shuttle in the enrichment might lower microbial diversity, natural environments are more likely to be dynamic and may contain many endogenous and exogenous organic and inorganic compounds that can function as electron shuttles for microbial Fe(III) reduction (6–8, 14, 15, 27, 39–45). When this is the case, our results suggest that microbial diversity could in fact be higher since we observe a unique microbial community with each different shuttle enrichment (Figure 4). Therefore, if we were to combine the unique community of each electron shuttle enrichment into a single community, we might expect the coexistence of many different taxa capable of using their preferred shuttle.

We also hypothesized that the presence of electron shuttles would improve the iron reduction rates. In this case, the iron reduction rates varied by shuttle. We observed an inhibition of iron reduction in the presence of the electron shuttle AQC compared to the no shuttle treatment and an increase in the iron reduction rate in the presence of AQDS (Fig 1D). These findings reinforce the idea that the presence of electron shuttles can have nuanced effects on biogeochemical cycling and microbial community function.

The nuanced effects that shuttles had on microbial activity cannot be explained by the redox potential of the shuttle. In fact, we observed a discrepancy between the present study and our earlier investigation on the correlation between redox potentials of the electron shuttles and Fe(III) reduction (40). In that work, the redox potential of the electron shuttle was directly correlated with the Fe(III) reduction rate by *Shewanella putrefaciens* CN32. Therefore, shuttles may have a clear redox-dependent effect on individual taxa but not a consistent effect on a mixed microbial community. Indeed, in contrast to O’Loughlin *et al*. (40), AQC had one of the lowest redox potential of the shuttles tested here (-0.247 V), but it exhibited the least iron reduction in this study whereas it led to the fastest iron reduction rate with *S. putrefaciens.* Low redox potential did not cause this inhibition of iron reduction as the shuttles AQDS (-0.225V) and riboflavin (-0.235V) have redox potentials close to that of AQC but are not deficient in iron reduction compared to the no shuttle treatments.

According to the end-point stoichiometric analysis, acetate consumption, iron reduction, and methane production were nearly the same across all of the shuttle treatments, with a notable absence of methanogenesis in the AQC treatment (Figure 1, Table 1). We hypothesize that the absence of methanogenesis in AQC treatments could be due to either a direct inhibition of methanogenesis by AQC or competition between microbial community members enriched under AQC. However, it is unlikely that AQC inhibited methanogenesis directly since *Methanosarcina* were enriched in many of the treatments in this study and *M. barkeri* has been shown to generate more methane when exposed to AQC (46). Therefore, we think it more likely that AQC enriched for a unique microbial community that either outcompeted, actively inhibited, or otherwise competitively excluded the methanogens early in the enrichments.

## CONCLUSIONS

Electron shuttle effects on Fe(III) reduction and methanogenesis were compound specific. Interestingly, AQC-amended experiments showed initially slower Fe(III) reduction and a complete inhibition of methanogenesis. Previous work under axenic conditions (*S. putrefaciens* CN32) indicated a robust relationship between an electron shuttle’s reduction potential and the rate of Fe(III) reduction, such that AQC>AQS>AQDS>AQZ>NQL>NS (40). However, in this study iron reduction rates were significantly slower in AQC compared to the rest of the treatments, suggesting that the reduction potential is not an effective predictor for the effectiveness of a putative electron shuttle in mixed microbial systems. Members of the Geobacteraceae and Geothermobacteraceae families dominated in the absence of added electron shuttles as well in certain electron shuttle treatments. The complete inhibition of methanogenesis by AQC highlights the possibility for electron shuttles to influence microbial processes in addition to those involving respiration with an insoluble terminal electron acceptor. Future work will focus on how mixtures of shuttles might change microbial diversity in Fe(III)-reducing environments and why certain shuttles might have inhibitory effects on microbial activity.

## MATERIALS AND METHODS

### Chemicals and Media

Natural sienna, an Fe-rich earth collected from ochre deposits in the Provence region of France consisting primarily of quartz and goethite (α-FeOOH) (47), was obtained from Earth Pigments Co. and used as received. NQJ, NQL, NQS, AQC, AQS, AQZ (see Table 1 for names of the shuttles), ferrozine, N-(2-hydroxyethyl) piperazine-N′-(2-ethanesulfonic acid) (HEPES), and piperazine-N,N′-bis(2-ethanesulfonic acid) (PIPES) were purchased from Sigma-Aldrich. AQDS was purchased from Fluka. Stock solutions (10 mM) of NQS, AQS, and AQDS were prepared in water and filter-sterilized (pore size = 0.22 µm). Due to their low aqueous solubility, 10 mM stock solutions of NQJ, NQL, and RIBO were prepared in methanol, and AQC and AQZ were prepared in acetone. A defined minimal medium consisting of 40 mM PIPES buffer, 40 mM HEPES buffer, 2 mM NH_4_Cl, 1 mM KCl, 2 mM CaCl_2_, 2 mM MgCl_2_, 20 µM phosphate, 20 mL of trace metal solution (40), and 20 mM sodium acetate, adjusted to pH 7.0, was sterilized by filtration through a 0.22 µm filter. A suspension of natural sienna was prepared by adding 100 g of natural sienna to 800 mL of water, which was then sonicated to ensure dispersion of the solids. The pH was adjusted to 6.8 and the suspension was autoclaved.

### Source of Inoculum

The inoculum for the bioreactors was prepared from sediment collected from a *Typha*-dominated wetland at Argonne National Laboratory in Lemont, IL, USA (41.710278, −87.985833). The top 5 cm of sediment were sealed in a coring device, then returned immediately to the laboratory and placed in an anoxic glove box (Coy Laboratory Products; N_2_:H_2_ = 95:5, [O_2(g)_] < 1 ppm). Core sediment was combined with overlying water from the wetland to create a slurry that was used to inoculate the bioreactors described below.

### Experimental System

Bioreactors were prepared in sterile 160-mL serum bottles. A volume of electron shuttle stock solution required to achieve a concentration of 100 µM in the bioreactor was added to the serum bottle. In the case of shuttles dissolved in methanol or acetone, the solvent was removed via evaporation under a stream of sterile air while the bottle was rotated, thus creating a film of the shuttle compound on the interior of the bottle. A 50 mL volume of the defined minimal medium and 50 mL of the natural sienna suspension was added to each bottle (the resulting systems contained 1 mmol of acetate and 20 mmol Fe(III)). The bottles were sparged with sterile argon (Ar), sealed with butyl rubber plug stoppers and aluminum crimp caps, and spiked with 1 mL (at STP) of xenon (Xe) as an internal standard for headspace analysis of CH_4_ and CO_2_. All systems were prepared in triplicate. The bottles were incubated at 30 °C in the dark with continuous mixing for a minimum of 24 h to allow for dissolution of the electron shuttle films. The enrichments were inoculated with 1 mL of sediment slurry and incubated at 30 °C in the dark with continuous mixing. A series of three experiments were conducted, and each was inoculated with a freshly prepared sediment slurry. Experiment 1 included bioreactors containing AQDS, NQL, and AQC; experiment 2 had AQZ, and AQS; and experiment 3 had NQS, NQJ, and RIBO. Each experiment included a no shuttle control, and all experimental systems were prepared in triplicate.

Sterile needles and syringes were used to collect samples of the headspace and suspension over time to monitor changes in the biogeochemistry and microbial communities in the bioreactors. A 200 µL sample of the headspace was collected to determine CH_4_ and CO_2_ concentrations. A 0.25 mL aliquot of suspension was added to 0.75 mL of anoxic 1 M HCl to measure Fe(II) concentration. A 1 mL aliquot of suspension was collected for measurement of acetate concentration and a 2 mL sample of suspension was collected and frozen at -80 °C for subsequent DNA extraction and sequencing.

### Analytical Methods

The reduction of Fe(III) in the bioreactors was monitored by the production of Fe(II) over time as described by Flynn et al. (47). Samples for acetate analysis were prepared by centrifuging 1 mL of suspension at 25,000 × *g* for 10 min and combining 0.5 mL of the supernatant with 0.5 mL of 10 mM H_2_SO_4_. The concentration of acetate was measured with an Agilent 1100 series HPLC equipped with a Bio-Rad Aminex HPX-87H ion exchange column (7.8 x 300 mm) eluted with 5 mM H_2_SO_4_ at a flow rate of 0.6 mL min^-1^ at 50 °C, with analyte detection at 210 nm. The concentrations of CH_4_ and CO_2_ were measured by analyzing 200 µL sample of headspace with an Agilent 7890A gas chromatograph as described by O’Loughlin et al. (48).

### DNA Extraction, Amplification, and Sequencing

DNA extraction was performed using the Qiagen PowerSoil DNA Isolation kit according to the manufacturer’s protocol. All DNA quantitation was performed using the Qubit assay (Invitrogen). PCR was used to amplify the V4 region of the 16S rRNA gene using a modified version of the universal 515F-806R primer pair in order to survey the bacterial and archaeal community in the extracted samples (49). Paired-end amplicons (2 × 151 base pairs) were then sequenced on the Illumina MiSeq platform using customized sequencing primers and procedures as described by Caporaso et al. (50). Samples were sequenced over three separate sequencing runs (Batches 1, 2, and 3), generating 6.91×10^6^ paired-end sequences for an average sequencing depth of (31,858 ± 20,617 sequences per sample). All extractions, amplification, and sequencing were performed at the Environmental Sample Preparation and Sequencing Facility (ESPSF) at Argonne National Laboratory (Lemont, IL, USA).

### Sequence analysis

Sequences were demultiplexed using idemp (51) and processed using the published tutorial pipeline of dada2 v1.14.1 on R v3.6.1(52). Sequencing runs were merged by dada2 after quality filtering and prior to chimera removal step. Merged sequences were assigned taxonomy using the SILVA v138 database (https://www.arb-silva.de/). Tree was built with phangorn (53) using aligned sequences from DECIPHER v. Processed sequences were analyzed using phyloseq v1.34.0 (54) and vegan v2.5-7 (REF) in R v4.0.4 .

### Data availability

Sequences were deposited to the NCBI sequence read archive (SRA) under BioProject PRJNA1064941 and accession numbers SAMN39446379 - SAMN39446765.

## ACKNOWLEDGEMENTS

Research under the Wetlands Hydrobiogeochemistry Scientific Focus Area (SFA) at Argonne National Laboratory was supported by the Environmental Systems Science Program, Office of Biological and Environmental Research (BER), Office of Science, U.S. Department of Energy (DOE), under contract DE-AC02-06CH11357. Argonne National Laboratory is a U.S. Department of Energy laboratory managed by UChicago Argonne, LLC. This work was supported in part by the U.S. DOE, Office of Science, Office of Workforce Development for Teachers and Scientists (WDTS) under the Science Undergraduate Laboratory Internship Program (SULI).

---

**Supplemental Figure S1.**
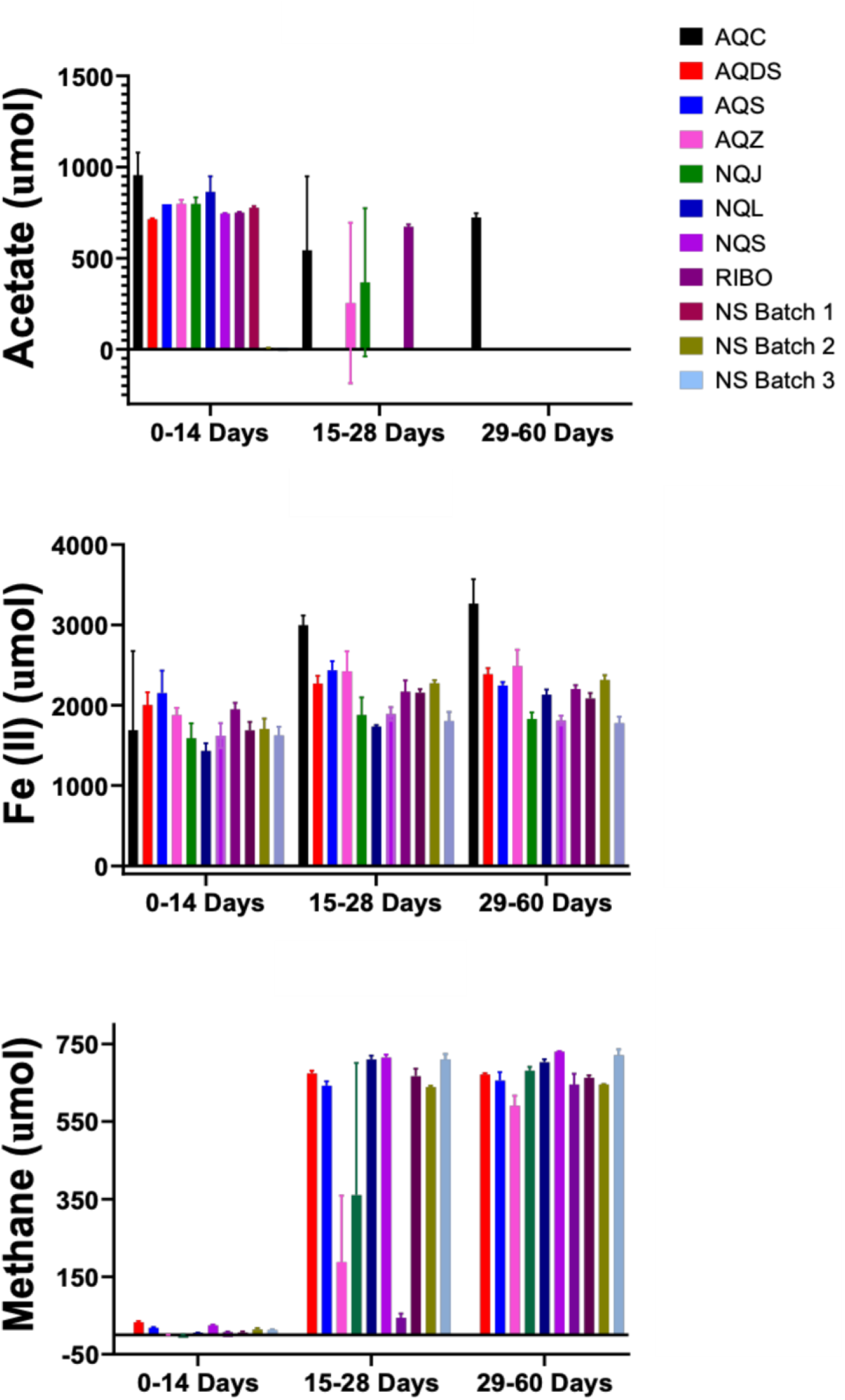
Iron(II) production, acetate consumption, or methane production in each experimental treatment visualized after three different phases of incubation - first two weeks of primarily iron reduction, middle two weeks transition from iron reduction to methane production, and remainder 32 days when acetate was completely consumed.

**Supplemental Figure S2.**
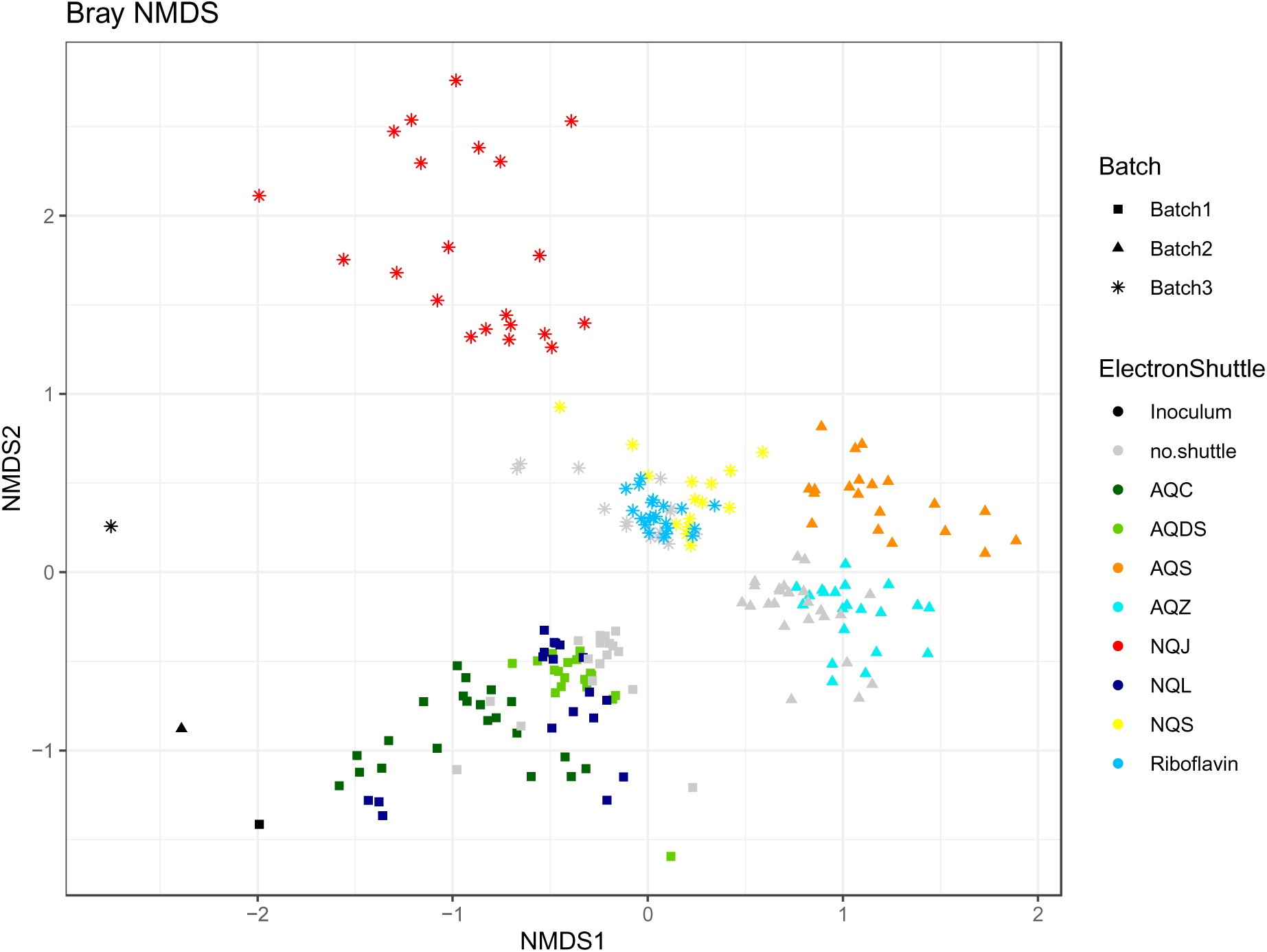
Relatedness of all communities measured as Bray-Curtis on an NMDS.

**Supplemental Figure S3.**
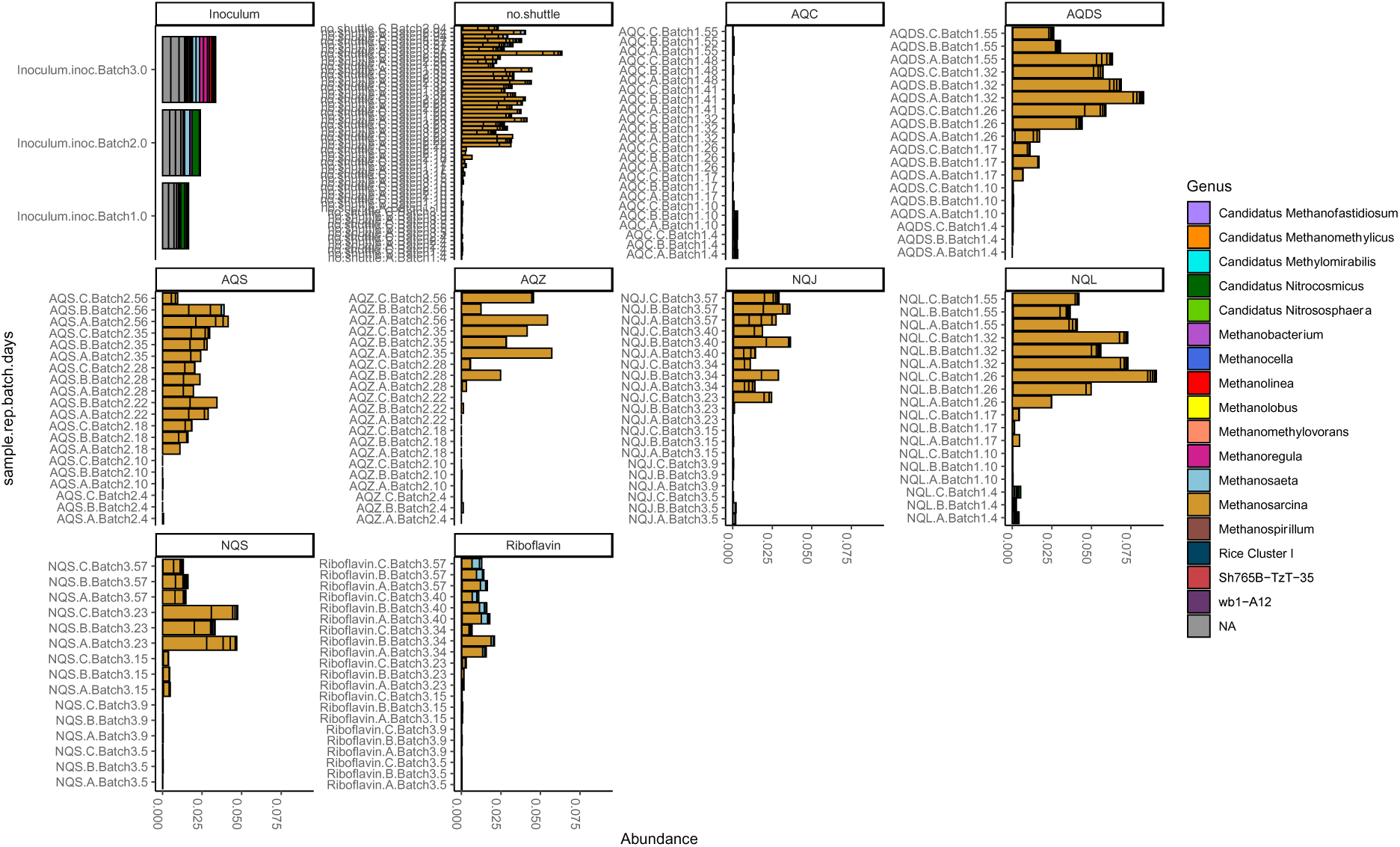
Relative abundance of ASVs belonging to Halobacterota, Euryarchaeota, Crenarchaeota, or Methylomirabilota phyla.

## Footnotes

1 Here we used the SILVA database(33) naming convention for all taxa but acknowledge the challenges associated with certain taxonomic assignments. The genus *Citrifermentans* in particular is controversial (34) and might be more appropriately named *Geomonas* (35), but we have left the nomenclature in place to allow for reproducibility and consistency.

